# Intra-genus metabolic diversity facilitates co-occurrence of multiple *Ferrovum* species at an acid mine drainage site

**DOI:** 10.1101/751859

**Authors:** Christen L. Grettenberger, Jeff R. Havig, Trinity L. Hamilton

**Affiliations:** Dept. of Earth and Planetary Sciences, University of California, Davis, CA USA 95616; Dept. of Earth and Environmental Sciences, University of Minnesota, Minneapolis, MN USA 55455; Dept. of Plant and Microbial Biology, University of Minnesota, Saint Paul, MN USA 55108; BioTechnology Institute, University of Minnesota, St. Paul, MN USA 55108

**Keywords:** *Ferrovum*, acid mine drainage, carbon fixation, iron oxidation, metagenome

## Abstract

**Background:** *Ferrovum* spp. are abundant in acid mine drainage sites globally where they play an important role in biogeochemical cycling. All known taxa in this genus are Fe(II) oxidizers. Thus, co-occurring members of the genus could be competitors within the same environment. However, we found multiple, co-occurring *Ferrovum* spp. in Cabin Branch, an acid mine drainage site in the Daniel Boone National Forest, KY.

**Results:** Here we describe the distribution of *Ferrovum* spp. within the Cabin Branch communities and metagenome assembled genomes (MAGs) of two new *Ferrovum* spp.. In contrast to previous studies, we recovered multiple 16S rRNA gene sequence variants suggesting the commonly used 97% cutoff may not be appropriate to differentiate *Ferrovum* spp. We also retrieved two nearly-complete *Ferrovum* spp. genomes from metagenomic data. The genomes of these taxa differ in several key ways relating to nutrient cycling, motility, and chemotaxis.

**Conclusions:** Previously reported *Ferrovum* genomes are also diverse with respect to these categories suggesting that the genus *Ferrovum* contains substantial metabolic diversity. This diversity likely explains how the members of this genus successfully co-occur in Cabin Branch and why *Ferrovum* spp. are abundant across geochemical gradients.

## 1. BACKGROUND

Iron-oxidizing bacteria are common in mine-impacted water and acid mine drainage (AMD) environments which are typically characterized by low pH and high concentrations of dissolved metals [1–3]. Betaproteobacteria of the genus *Ferrovum* play important roles in biogeochemical cycling in AMD environments including carbon fixation and rapid oxidation of iron [4–9] and could be of value in bioremediation [10]. However, *Ferrovum* spp. are challenging and labor-intensive to culture and isolate because they often co-occur with heterotrophic *Acidiphilium* spp. or other Fe(II) oxidizers [7, 11, 12]. Indeed, *Ferrovum myxofaciens* strain P3 is the only *Ferrovum* spp. in culture to date, highlighting the value and need for characterization of this enigmatic genus via molecular techniques.

*Ferrovum* spp. occur in AMD environments with diverse geochemistry. *Ferrovum* spp. are abundant at sites with pH ranging from less than 3 to 7 [4–6, 13, 14] and with iron concentrations ranging from 2 μM [5] to 71 mM [7]. High-throughput sequencing and ‘omics techniques have aided in characterizing the metabolic potential of *Ferrovum* via non-culture-based techniques [15–18] across these large gradients of Fe and pH. These studies indicate that *Ferrovum* is a diverse genus composed of six clades (Groups I - VI) of closely related species and strains [18]. Still, only four *Ferrvoum* genomes are publicly available and these are from *Ferrovum* Groups I and IV. No genomes have been reported for Groups, II, III, IV, or V. The published genomes and culture studies suggest that *Ferrovum* spp. are autotrophs that pair Fe(II) oxidation with carbon fixation [7, 15, 16, 18, 19]. Some members of the genus also fix nitrogen and are therefore important to nutrient cycling [7, 19]. However, sequenced *Ferrovum* genomes have variable genomes especially with genes associated with motility, chemotaxis, biofilm formation, and nitrogen metabolism [15]. Furthermore, *Ferrovum* dominated communities exhibit morphological differences, including forming large streamers [5, 7, 14] or low extracellular polymeric substances [4] though it can be difficult to identify EPS in low pH environments [16]. These differences may help explain how *Ferrovum* is able to occupy a large range of pH and ferrous iron concentrations in AMD environments, However, the potential genetic and metabolic diversity within the genus is under-represented because there are no cultured representatives or genomes available for several *Ferrovum* clades.

In our previous studies of Cabin Branch, we reported the predominance of a single operational taxonomic unit (OTU; 97% similarity) in our 16S rRNA amplicon data most closely related to *F. myxofaciens* at an AMD site in the Daniel Boone National Forest in southern Kentucky (Figure 1). This taxon was abundant in diverse morphotypes including filament, floc, and ‘brain’ or ‘spongy’ mat biofilms over a pH range of 2.1 to 2.4 and Fe(II) concentrations of 448 to 5200 μmol/L [5]. Based on these data, we employed a cultivation-independent approach to characterize the genetic diversity and functional potential of this taxon using 16S rRNA and by examining the gene content of metagenome assembled genomes (MAGs) from the site.

**Figure 1.**
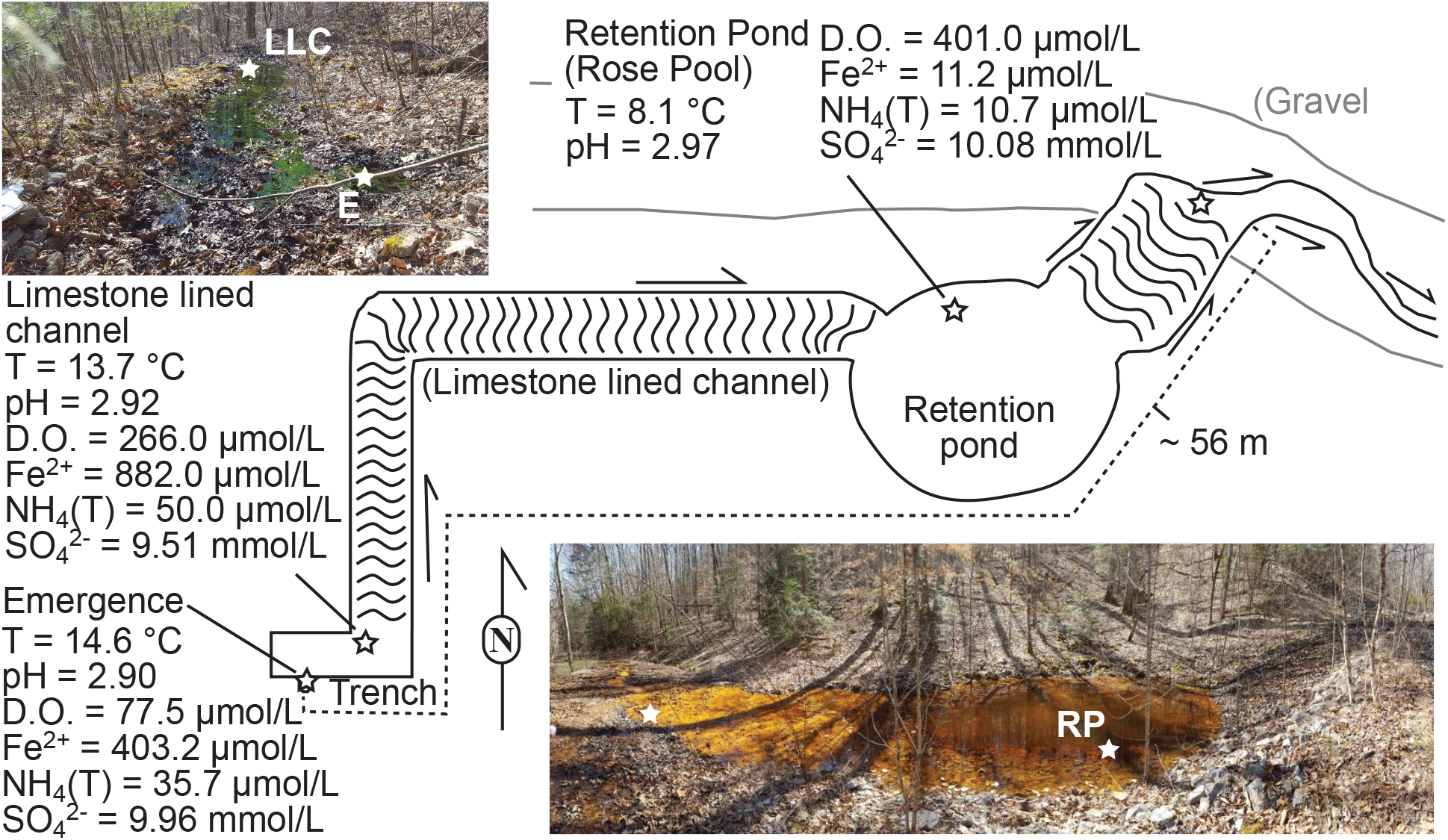
Site location.

## 2. RESULTS

### 2.1 Geochemistry

pH values were acidic, ranging from 2.90 to 2.97 from the source to Rose Pool. Temperatures were highest at the source (14.6 °C) and decreased down the outflow channel to 8.1 °C at the Rose Pool, consistent with warm groundwater cooling after emerging due to relatively cool ambient temperatures (~ 0 °C at the time of sampling). Conductivity increased from 940 μS/cm at the emergence to 2063 μS/cm at Rose Pool. Dissolved oxygen increased from a low of 77.5 μmol/L at the emergence sampling site to a high of 401 μmol/L (close to saturation) at Rose Pool. Anion concentrations in Cabin Branch were dominated by sulfate (9.51 to 10.08 mmol/L), with lower concentration of chloride (166 to 220 μmol/L). Cation concentrations were highest at the emergence and nearby first outflow sites, and lowest at the Rose Pool, including calcium (1.20 to 0.39 mmol/L), magnesium (0.69 to 0.20 mmol/L), potassium (0.18 to 0.03 mmol/L), and sodium (0.12 to 0.03 mmol/L).

Dissolved inorganic carbon (DIC, predominantly present as dissolved CO_2_ due to the low pH) concentration were highest at the emergence (1.67 mmol/L) and decreased down the outflow channel to 0.30 mmol/L at the Rose Pool. DIC δ^13^C values were more negative at the source (δ^13^C = −15.96 ‰) and became more positive down the outflow channel with the most positive values at the Rose Pool (δ^13^C = −11.78 ‰), consistent with preferential loss of ^12^CO_2_ through volatilization and uptake by autotrophs. Dissolved organic carbon concentration was consistently low at all sites (36.8 to 44.2 μmol/L) and had similar δ^13^C values (−22.50 to −23.95 ‰).

Potential nutrients measured included NH_4_(T), P, Mn, Fe(II), and Fe_(total)_. NH_4_(T) concentration was highest at the first outflow site (50.0 μmol/L) and lowest at Rose Pool (10.7 μmol/L). P concentration was highest at the emergence (9.55 μmol/L) and decreased to below detection limits at the Rose Pool. Mn concentration decreased from the highest value at the emergence (76.84 μmol/L) to the lowest at the Rose Pool (40.23 μmol/L). Total iron (Fe_total_) exhibited a similar trend as Mn and P, with the highest concentration at the emergence (4.68 mmol/L) and the lowest at the Rose Pool in the outflow (0.25 mmol/L), though Fe(II) had concentrations significantly lower than Fe_total_ with the highest concentration at the outflow site downstream of the emergence (0.88 mmol/L) and the lowest at the Rose Pool (11.2 μmol/L). The discrepancy in the Fe(II) data may be due to the colorimetric tests which are optimized for 20 °C. whereas Fe_total_ is measured via ICP-MS or ICP-OES with samples that are acidified with concentrated HNO_3_ to keep metals in solution. All aqueous geochemistry data are reported in Table 1.

**Table 1.** Geochemical and physical analytical results for Cabin Branch samples.

| Parameter | Location:<br>(Units) | Emergence |  | Outflow 1 |  | Outflow 2 |  | Outflow 3 |  |
| --- | --- | --- | --- | --- | --- | --- | --- | --- | --- |
|  |  | Value | S.D. | Value | S.D. | Value | S.D. | Value | S.D. |
| pH |  | 2.90 |  | 2.92 |  | 2.97 |  | 3.04 |  |
| mV |  | 235 |  | 232 |  | 224 |  | 220 |  |
| Temperature | (°C) | 14.6 |  | 13.7 |  | 8.1 |  | 6.9 |  |
| Conductivity | (μS/cm) | 940 |  | 1853 |  | 2063 |  | 966 |  |
| D.O. | (μmol/L) | 77.5 |  | 266 |  | 401 |  | 276 |  |
| Fe<sup>2+</sup> | (μmol/L) | 403.2 |  | 882.0 |  | 11.16 |  | bdl |  |
| NH<sub>4</sub>(T) | (μmol/L) | 35.7 |  | 50.0 |  | 10.7 |  | 35.0 |  |
| DIC | (mmol/L) | 1.67 |  | 0.48 |  | 0.30 |  | 0.18 |  |
| δ<sup>13</sup>C<sub>VPDB</sub> | (‰) | -15.96 | 0.08 | -13.03 | 0.08 | -11.79 | 0.08 | -15.65 | 0.08 |
| DOC | (μmol/L) | 40.51 | 0.90 | 44.19 | 3.27 | 36.82 | 6.42 | 42.81 | 7.64 |
| δ<sup>13</sup>C<sub>VPDB</sub> | (‰) | -22.50 | 1.53 | -23.09 | 0.84 | -23.95 | 0.29 | -25.41 | 1.30 |
| Cl | (μmol/L) | 175.58 | 1.10 | 220.04 | 61.99 | 166.13 | 0.44 | 167.97 | 0.95 |
| SO<sub>4</sub> | (mmol/L) | 9.96 | 0.16 | 9.51 | 0.14 | 10.08 | 0.06 | 4.32 | 0.10 |
| P | (μmol/L) | 9.55 | 0.49 | 4.49 | 0.35 | bdl |  | 0.58 | 0.46 |
| Na | (mmol/L) | 0.12 | 0.00 | 0.12 | 0.03 | 0.03 | 0.00 | 0.07 | 0.00 |
| K | (mmol/L) | 0.17 | 0.00 | 0.18 | 0.04 | 0.03 | 0.00 | 0.09 | 0.00 |
| Ca | (mmol/L) | 1.121 | 0.028 | 1.198 | 0.256 | 0.390 | 0.007 | 0.850 | 0.018 |
| Mg | (mmol/L) | 0.666 | 0.012 | 0.686 | 0.156 | 0.201 | 0.001 | 0.404 | 0.002 |
| Al | (μmol/L) | *1755* | *125* | 1201 | 0.26 | 187.4 | 1.2 | 388.4 | 1.6 |
| Si | (μmol/L) | *1528* | *35* | *1454* | *66* | nd |  | *706* | *45* |
| Mn | (μmol/L) | 76.84 | 0.63 | 43.19 | 0.13 | 40.23 | 0.28 | 24.78 | 0.66 |
| Fe<sub>Total</sub> | (μmol/L) | *4684* | *22* | *4154* | *50* | 249.6 | 1.8 | *1154* | *46* |

### 2.2 Inorganic carbon assimilation

Autotrophic incorporation of added ^13^CO_2_ was quantified in mesocosm experiments using treatments carried out under either light or dark (wrapped in aluminum foil) to characterize photoautotrophic and chemoautotrophic uptake, respectively. At the emergence light treatments of full biofilms (green and white biomass) returned average C-uptake rates of 32.547 (± 7.949) and 21.691 (± 6.36) μg C uptake/g C_biomass_/hr after one- and two-hour incubations, respectively. Dark treatments of the green biomass returned averages of 1.632 (± 0.454) μg C uptake/g C_biomass_/hr after one hour and 0.746 (± 0.174) μg C uptake/g C_biomass_/hr after two hours, while dark treatments of the white biomass returned averages of 0.305 (± 0.403) μg C uptake/g C_biomass_/hr after one hour and 0.662 (± 0.240) μg C uptake/g C_biomass_/hr after two hours. Biofilms with both green and white biomass together from the limestone-lined channel immediately downstream of the emergence returned average C-uptake rates of 35.947 (± 5.516) and 15.085 (± 0.106) μg C uptake/g C_biomass_/hr for light treatments of one and two hours respectively and 2.586 (± 0.148) and 1.245 (± 0.035) μg C uptake/g C_biomass_/hr for dark treatments after one and two hours. Rose Pool sediment incubations returned average C-uptake rates of 1.372 (± 0.061) μg C uptake/g C_biomass_/hr for light treatments of two hours and 0.066 (± 0.132) μg C uptake/g C_biomass_/hr for dark treatments of two hours. All C-uptake results are reported in Supplementary Table S1.

### 2.3 16S rRNA

Cabin Branch contains nine denoised sequence variants (DSVs) closely related to *Ferrovum myxofaciens* (Figure 2). Four DSVs are present in the emergence sample, five in the limestone lined channel, and eight in Rose Pool (Figure 2, 3). DSVs affiliated with *Ferrovum* are most abundant in the limestone lined channel where they compose 32.8% of the community and least abundant in the Rose Pool where they compose 5.0% of the community. *Ferrovum* DSVs compose 29.6% of the emergence community. The abundance of each DSV varies between samples. DSVs 4, 10, 12, and 81 are the most abundant *Ferrovum* DSVs in the Emergence community, 4, 10, 12, 34, and 59 in the limestone lined channel, and DSVs 56 and 81 in the Rose Pool (Figure 3). The DSVs are phylogenetically placed within *Ferrovum* groups I, II, III and between Groups IV and V (Figure 2).

**Figure 2.**
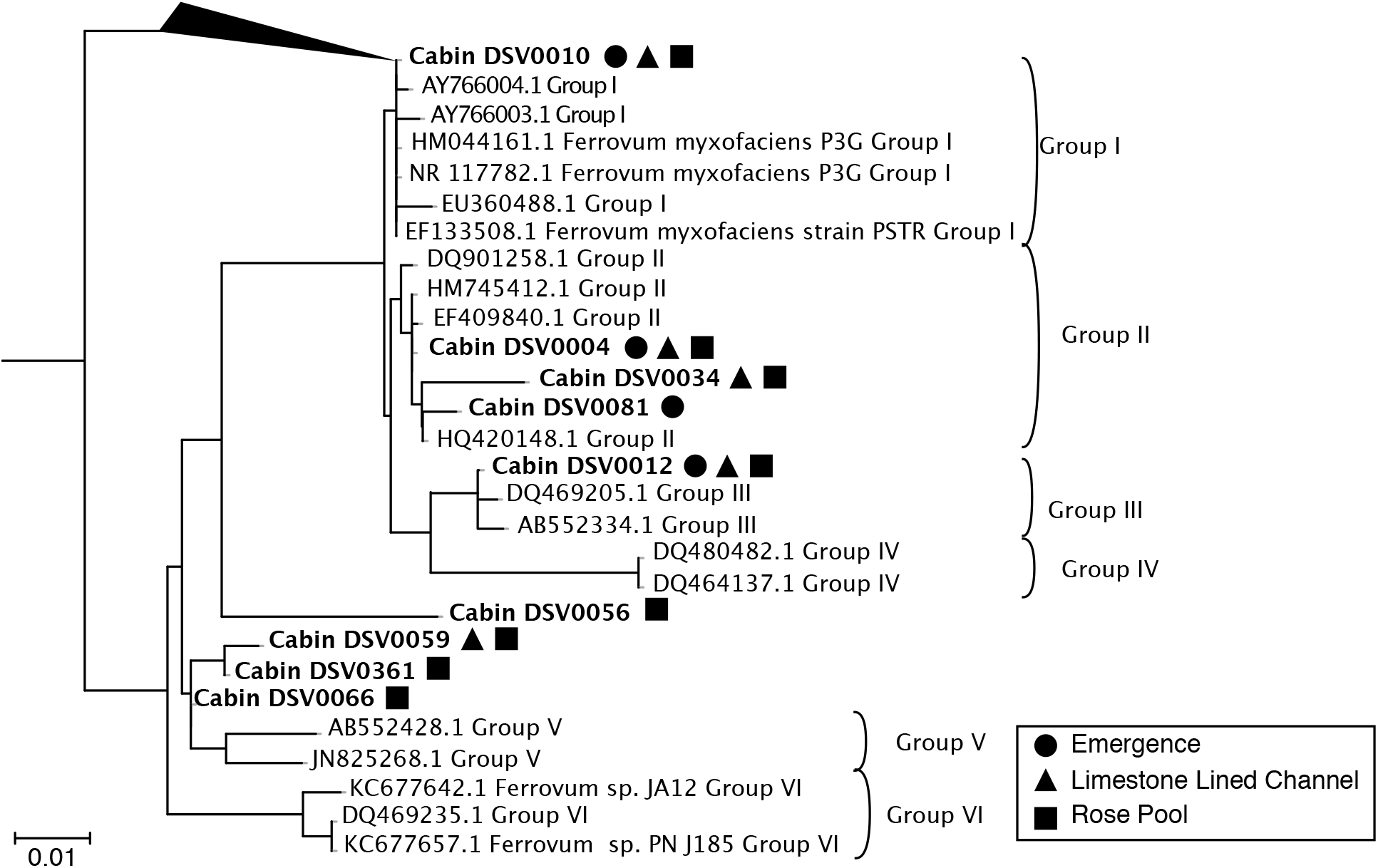
16S rRNA gene tree of *Ferrovum* spp. Group nomenclature from [18]. DSVs found in the emergence are indicated by circles, triangles indicate those found in the limestone lined channel, and squares those found in the Rose Pool.

**Figure 3.**
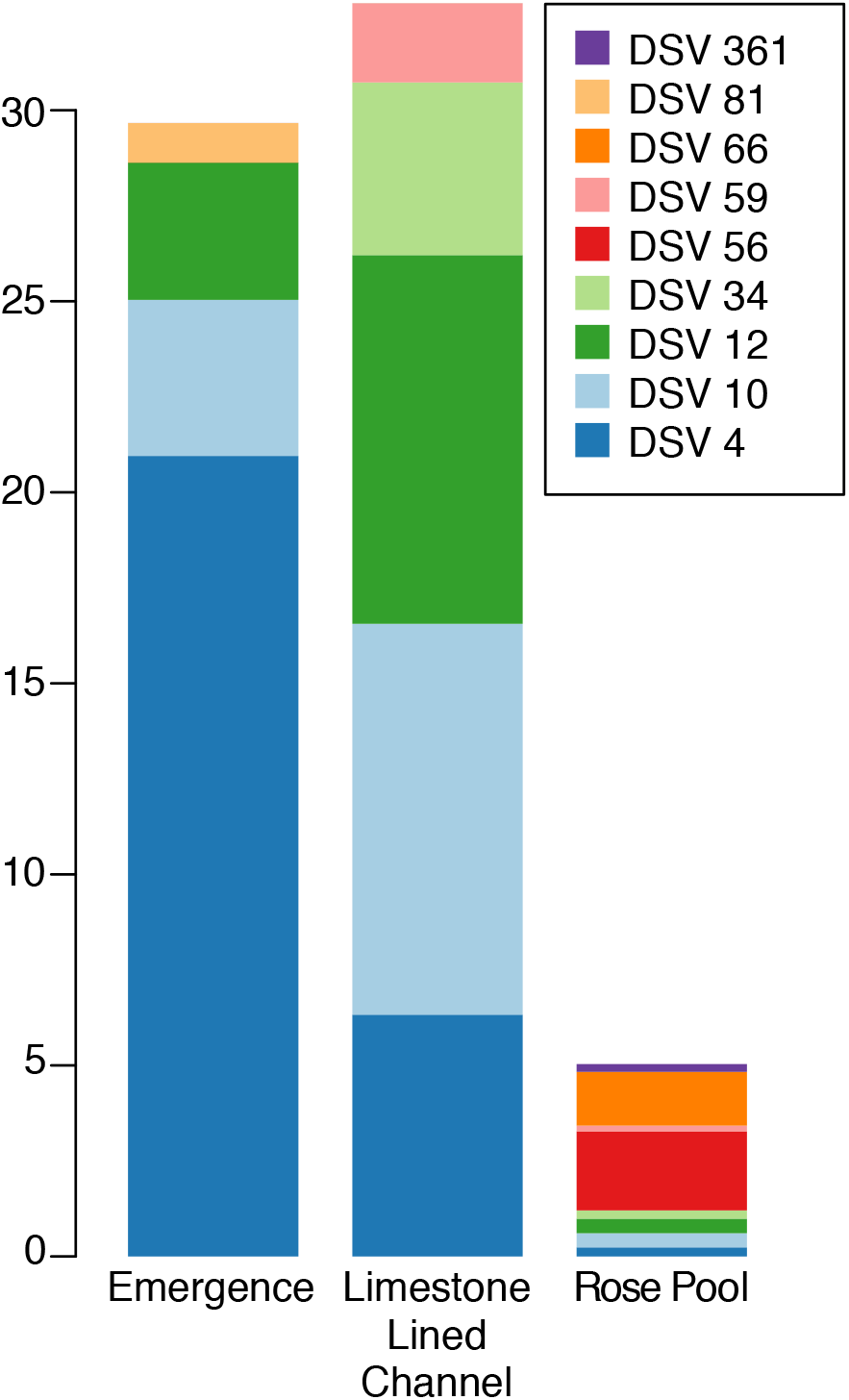
Relative abundance of *Ferrovum* DSVs in each of the sampled communities.

### 2.4 Metagenomics

Fourteen bins were retrieved that were phylogenetically placed within the genus *Ferrovum*. The resulting MAGs ranged from low- to high-quality drafts based on quality and contamination, but lack 16S rRNA gene sequences [20]. Five of these bins were retrieved from individually assembled metagenomes and 9 from the combined assembly. Based on ANI, each of the *Ferrovum* bins from the individual assemblies was also present in the combined assembly and one bin was represented in the emergence, terrace, and combined assemblies (Supplementary Table S2). For MAGs that were present in more than one assembly, the highest quality bin from that MAG was used for further analysis (Table 2). Nine unique *Ferrovum* MAGs were present in the dataset. These MAGs shared 68.5% to 88.5% ANI with each other and between 68.6% and 99.4% ANI with published *Ferrovum* genomes (Supplementary Table S2). Based on a concatenated tree of ribosomal protein genes, eight of the nine MAGs occupy the phylogenetic space between the Group I *Ferrovum* sp. JA12 and sp. PN-J185 and the Group IV *Ferrovum* sp. Z-31 and the type strain (Figure 4). Two MAGs, MAG-4 and MAG-7 are classified as high-quality drafts based on the criteria described by Bowers et al., [20] and are described in more detail below.

**Table 2.** Metagenome assembled genomes. When a taxon was found in multiple assemblies, the MAG with the highest completeness and lowest contamination was used. The MAGs used for further analysis are shown in black text whereas those not used are shown in grey text. The MAGs used in the comparative analysis are highlighted by in grey.

| Name | Bin ID | Completeness | Contamination | Heterogeneity | Protein Coding Sequences | Annotated Sequences |
| --- | --- | --- | --- | --- | --- | --- |
| MAG 1 | Combined_89 | 20.69 | 0 | 0 |  |  |
| MAG 2 | Combined_19 | 44.72 | 0.42 | 66.67 |  |  |
| MAG 3 | Rose Pool 43 | 68.75 | 4.13 | 18.18 |  |  |
|  | Combined_70 | 58.28 | 5.17 | 40 |  |  |
| MAG 4 | Combined_34 | 95.76 | 0.5 | 0 | 2250 | 1466 |
| MAG 5 | Combined_31 | 43.89 | 0 | 0 |  |  |
|  | Rose Pool 56 | 36.51 | 0 | 0 |  |  |
| MAG 6 | Combined_120 | 42.16 | 1.72 | 0 |  |  |
| MAG 7 | Limestone Lined Terr. 25 | 86.7 | 0.63 | 100 | 1764 | 1158 |
|  | Emergence 33 | 72.59 | 0 | 0 |  |  |
|  | Combined_101 | 70.86 | 0 | 0 |  |  |
| MAG 8 | Combined_72 | 86.37 | 33.74 | 80.26 |  |  |
|  | Emergence 23 | 83.17 | 54.31 | 81.58 |  |  |
| MAG 9 | Combined_5 | 30.17 | 0 | 0 |  |  |

**Figure 4.**
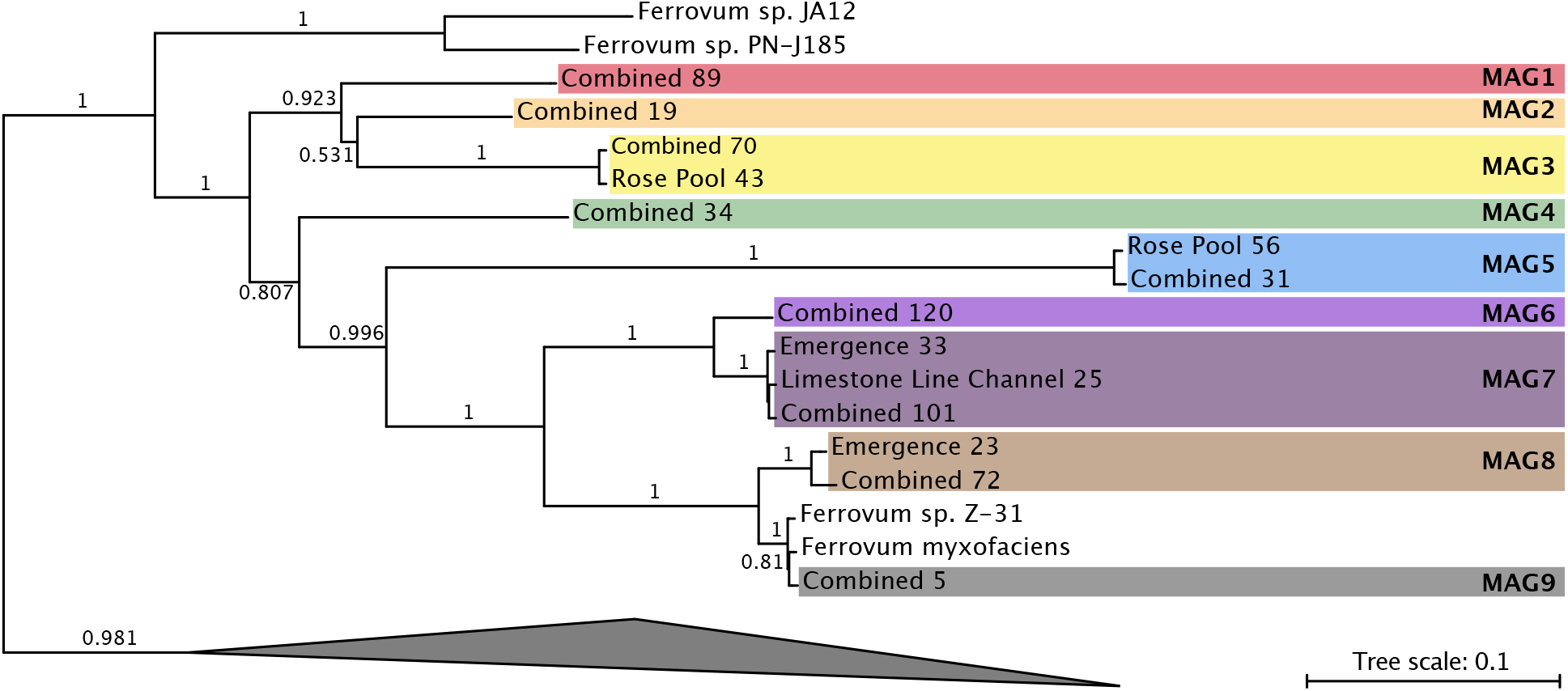
Concatenated gene tree containing all *Ferrovum* bins retrieved from the metagenomic datasets. Colored rectangles indicate the bins that belong to a given MAG. Bootstrap values (based on 100 bootstrap samplings) are shown for each node where bootstrap support is > 50%.

### 2.5 MAG-4

MAG-4 is represented by a single bin which was retrieved from the combined assembly and is the most complete MAG recovered in this study. It is 95.7% complete with 0.4% contamination. MAG-4 contains 2250 gene coding sequences, 1466 of which were annotated by GhostKoala (Table 2). MAG-4 shares 69 −74% ANI with the other MAGs and the published *Ferrovum* genomes (Supplementary Table S2). The MAG is phylogenetically positioned between the Group I *Ferrovum* spp. JA12 and PN-J185 and the Group VI type strain and sp. Z-31 (Figure 4)

#### 2.5.1 Energy Metabolism

MAG-4 contains three homologs of the high molecular weight, Cyc-2-like protein previously found in *Ferrovum* spp. (Figures 5, 6). This protein is thought to facilitate the oxidation of Fe(II) to Fe(III)[16]. The MAG also contains homologs of the genes necessary for oxidative phosphorylation using a B/A type NADH dehyrodgenase (*nuoA-nuoN*) and succinate dehydrogenase (*sdhA-D*), a E/B/A type cytochrome *c* reductase, *cbb*3 and *bd* type terminal oxidases, and an F-type ATPase (Figure 6).

**Figure 5.**
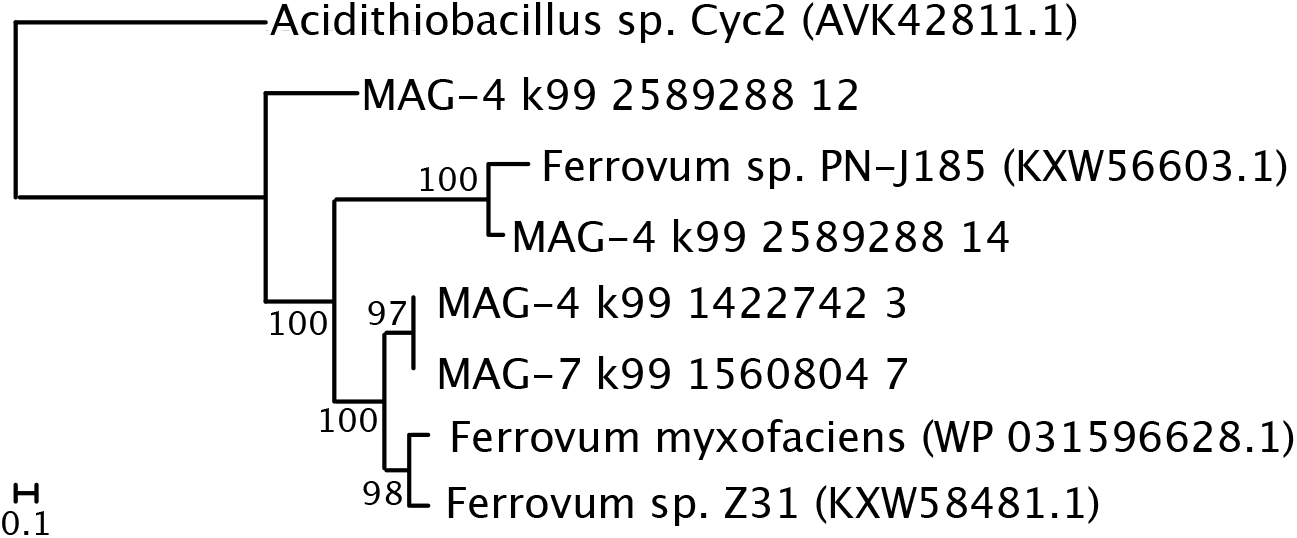
Maximum likelihood phylogeny phylogeny of Cyc-2 like gene found in published *Ferrovum* genomes and the MAGs retrieved in this study. Accession numbers are provided in parentheses. Numbers represent bootstrap support values based on 100 bootstrap samplings.

**Figure 6.**
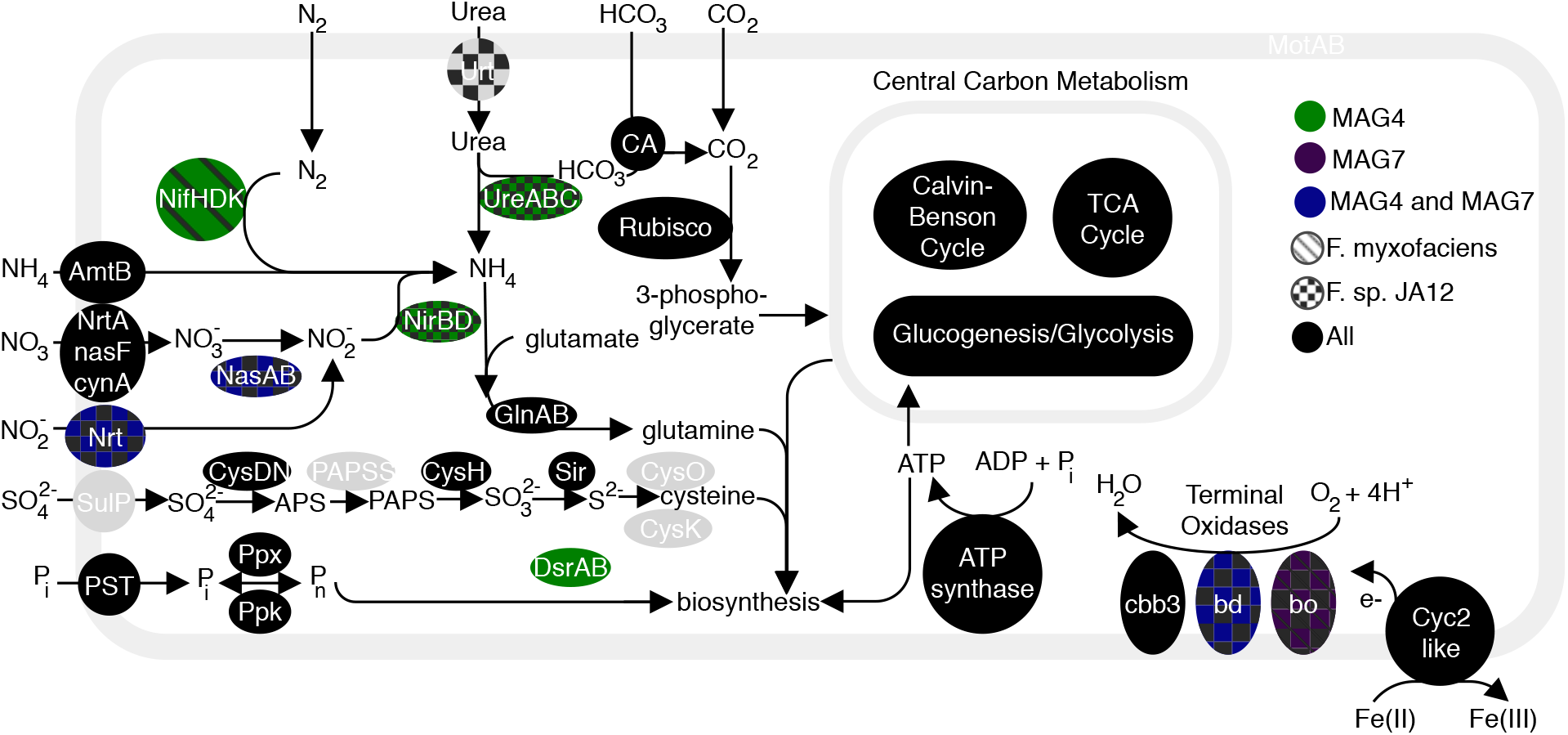
Predicted assimilation of C, S, P, and N in *Ferrovum* spp. Proteins are color coded based on their presence or absence in each genome. Modeled after [18].

#### 2.5.2 Carbohydrate Metabolism

MAG-4 encodes the enzymes necessary to fix carbon via the Calvin-Benson cycle except the enzyme needed to form ribose 5-phosphate from sedoheptulose-7 phosphate. The MAG also contains the genes for a nearly-complete TCA cycle. Only the gene for malate dehydrogenase (*mgo*) is missing, which catalyzes the oxidation of malate to oxaloacetate. It contains a nearly complete suite of genes for glycolysis but is missing the genes necessary to transform fructose-6-phosphate to fructose-1,6-bisphosphate (phosphofructokinase-1) and from glycerate-3-phosphate to glycerate-2-phosphate (triose-phosphate isomerase).

#### 2.5.3 Nutrient Acquisition

MAG-4 encodes the structural proteins necessary for nitrogen fixation (NifHDK) and denitrification (NasAB, NirBD), but lacks the genes necessary to import inorganic nitrogen in to the cell (Figure 6). It is likely that The MAG contains many of the genes necessary for assimilatory sulfate reduction but is missing PAPSS which reduces ammonium persulfate (APS) to 3-phosphoadenosine-5-phosphosulfate (PAPS). It also contains homologs of DsrA and DsrB which can be used to reduce sulfite to sulfide or to oxidize sulfide to sulfite (Figure 6). The MAG also contains the genes necessary to import inorganic phosphate and use it in biosynthetic pathways (*PST, ppx, ppk*).

#### 2.5.4 Motility

MAG-4 contains all the genes necessary to construct flagella except *motXY* which encode proteins are involved in flagellar rotation and *fliJT*. The MAG also contains the genes necessary for chemotaxis (*cheABCD*, *cheR*, and *cheVWZ*)

### 2.6 MAG-7

MAG-7 is represented by bins retrieved from the emergence, limestone lined channel, and co-assembled metagenomes (Table 1). The highest quality bin is from the limestone lined terrace and is 86.7% complete with 0.63% contamination. The other two bins are 72.6% and 70.9% complete with 0% contamination. Because the most complete bin is only 86.7% complete, when genes were absent from that bin, we searched the other two less complete bins

#### 2.6.1 Energy Metabolism

MAG-7 contains one homolog to the high-molecular weight Cyc2 type protein found in other *Ferrovum* taxa (Figure 5, 6). The Cyc2 sequence is phylogenetically placed as a sister group to those from the type strain and *F.* sp. Z31 (Figure 5). MAG-7 also contains homologs of the genes necessary for oxidative phosphorylation with a B/A type NADH dehydrogenase (*nuoA-nuoN*) and succinate dehydrogenase (*sdhA-D*), an E/B/A type cytochrome *c* reductase, *bo*, *bd*, and *cbb*3 type terminal oxidases, and an F-type ATP-ase.

#### 2.6.2 Nutrient Acquisition

The MAG encodes the genes necessary for reducing nitrate to nitrite (*nasAB*), but not those to further reduce the nitrate to ammonia (*nirBD*; Figure 6). The MAG also contains the genes necessary to import inorganic phosphate and use it in biosynthetic pathways (*PST, ppx, ppk*). MAG-7 does not encode protein necessary for nitrogen fixation and lacks the genes necessary to transform urea to ammonia.

#### 2.6.3 Carbohydrate Metabolism

MAG-7 contains most of the genes necessary for carbon fixation via the Calvin-Benson cycle but like MAG-4 does not encode the enzyme needed to form ribose 5-phosphate from sedoheptulose-7 phosphate. The MAG encodes the proteins necessary for a complete TCA cycle. It contains a nearly complete suite of genes for glycolysis but is missing the protein necessary to transform glycerate-3-phosphate to glycerate-2-phosphate.

#### 2.6.4 Motility

MAG-7 contains all the genes necessary to synthesize a functional flagellum except *motX* and *motY* which are also missing from MAG-4. The MAG also contains genes involved in bacterial chemotaxis including *mcp*, *aer*, *tar, cheABCD*, *cheR*, and *cheVY*, but it does not appear to contain *cheW* and *cheZ* which are present in MAG-4.

## 3. DISCUSSION

### 3.1 Co-occurrence of *Ferrovum* spp. in Cabin Branch Communities

Co-occurring strains of the same species have been found in a variety of environments including cyanobacterial blooms (Willis et al., 2018), and solar saltern/crystallization ponds [21, 22]. In some communities, successful co-occurrence results from genomic variation or differential gene expression between the strains of the co-occurring species. For example, six co-occurring strains of the Epsilonproteobacteria *Lebetimonas* acquired new functional genes via lateral gene transfer in sea mounts in the Mariana Arc [23]. Similarly, strains of *Salinabacter ruber* contain hypervariable regions in their genome which are associated with differences in surface properties [21]. Co-occurring strains of *S. ruber* also express different metabolite pools [22]. These differences likely allow co-occurring strains to avoid competing within the environment [21, 23]. Like these communities, the *Ferrovum* spp. at Cabin Branch may be able to co-exist due to variations in their genomes or in gene expression. All known *Ferrovum* spp. are obligate, Fe(II) oxidizing autotrophs that use O_2_ as a terminal electron acceptor [7, 16, 17]. Therefore, co-occurring *Ferrovum* spp. would presumably compete for common resources potentially including Fe(II) and oxygen as well as essential nutrients (e.g., N and P) and trace elements. At Cabin Branch, Fe(II) is present in mmol/L amounts constantly supplied by mine-impacted sources and is unlikely to be limiting. Dissolved oxygen is also present although the concentration is lower at the emergence and increases downstream. Ammonia and phosphate were detected at all sites. While we cannot rule out the effects of micronutrients, in the presence of presumably replete resources, gene content differences could explain co-existence of *Ferrovum* populations at Cabin Branch.

*Ferrovum* spp. compose over 25% of the bacterial community in the emergence and limestone lined channel communities and 5% of the Rose Pool community, based on the 16S rRNA amplicon libraries. The abundance of *Ferrovum* spp. correlates with the rate of non-photosynthetic inorganic carbon assimilation which suggests that these populations are important primary producers in the environment (Figure 7). Our previous work reported a single *Ferrovum* OTU in Cabin Branch. However, that analysis relied on a definition of 97% identity in 16S rRNA sequences [5] which is the common cutoff for delineating species based on 16S rRNA gene sequence identity level, but likely underestimates bacterial diversity [24]. Currently, methods using unique sequences are becoming more common and may be more robust [25, 26]. Using a DSV approach, we recovered nine unique *Ferrovum* taxa: four *Ferrovum* DSVs co-occur in the emergence community, five in the limestone lined channel, and eight in the Rose Pool sediments (Figures 2, 3). Genome-based studies using average nucleotide identity (ANI) and multilocus phylogenetic analysis are more robust than 16S rRNA-based analyses and are the best practice for assigning species affiliations in genomes [27]. Metagenomic data from Cabin Branch support the presence of multiple *Ferrovum* spp. at each site, suggesting that the diversity of *Ferrovum* spp. at AMD sites globally may be underestimated because studies have traditionally used an OTU approach [4, 5].

**Figure 7.**
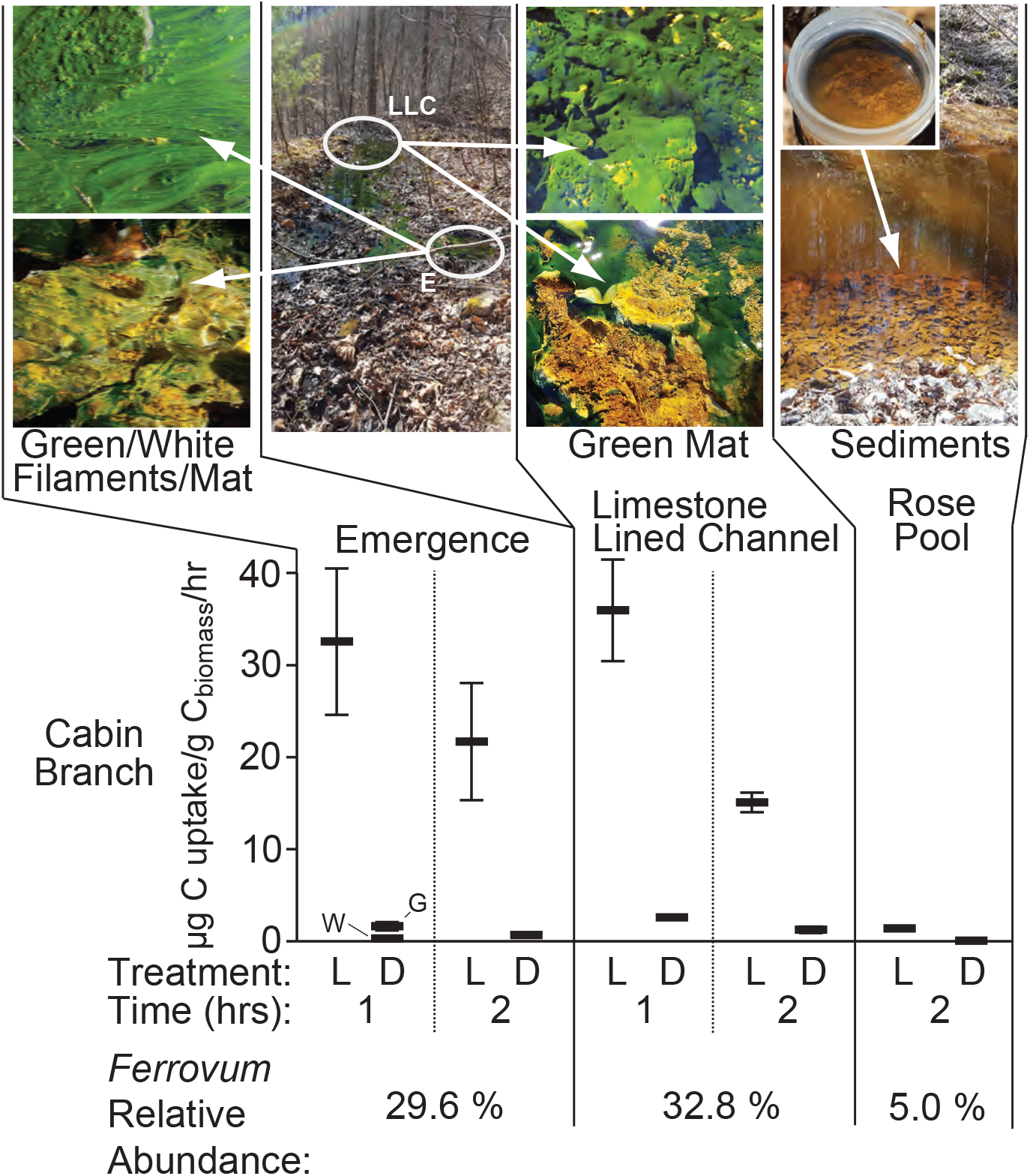
Images and carbon uptake rates for Cabin Branch emergence (E), limestone lined channel (LLC), and Rose Pool retention pond sites. L = light treatment, D = dark treatment (wrapped in aluminum foil), W = white filaments/mat, G = green filaments/mat.

To co-occur successfully, the strains may partition the niche to use different resources as is seen in closely-related taxa or intraspecific competitors in eukaryotic systems [28–30]. If phylogenetic distance correlates with metabolic diversity, we might expect co-occurring strains of *Ferrovum* would be distantly related. Indeed, those that co-occur in the Cabin Branch communities span the Group I, II, and III *Ferrovum* clades and the limestone lined channel and Rose Pool also include DSVs that are not placed within a clade (Figure 2). However, some co-occurring *Ferrovum* spp. are closely related to one another. Multiple DSVs from the same *Ferrovum* group are found in the same microbial community. For example, two Group II DSVs are found in all three communities sampled and the Rose Pool community contains three closely related DSVs phylogenetically placed basally to the Group V *Ferrovum* (Figure 2). Therefore, phylogenetic distance cannot be used as a proxy for the features that allow strains to co-exist. However, we can examine how genome content may allow co-occurring taxa to partition resources and avoid competition.

### 3.2 Relating two new *Ferrovum spp.* (MAG-4 and MAG-7) to published species

Here, we compared our *Ferrovum* MAGs to two published *Ferrovum* species from *Ferrovum* Groups I (*F. myxofaciens* sp. P3G) and IV (*Ferrovum* JA12) to explore if genomic variability could contribute to the co-occurrence of these taxa. Our two *Ferrovum* MAGs and the Group I and Group IV genomes share numerous features, but differ in key ways relating to energy metabolism, nitrogen cycling, and motility.

#### 3.2.1 Energy metabolism

All cultured *Ferrovum* spp. can grow autotrophically via iron oxidation [7, 15, 16, 18, 19] and we predict that the two MAGs presented here share that ability. Each genome contains high molecular weight, Cyc2-like proteins found in other *Ferrovum* taxa [16]. Cyc2-like proteins are found in many acidophilic and neutrophilic Fe(II) oxidizers and are highly expressed in their metatranscriptomes [31]. Therefore, this protein likely plays a role in Fe(II) oxidation. In autotrophic Fe(II) oxidizers like *Ferrovum spp.*, *Acidithiobacillus ferrooxidans*, and *Gallionella ferruginea*, Fe(II) oxidation is paired with carbon fixation. The MAGs encode a nearly full suite of genes for carbon fixation via the CBB pathway but differ in the types of terminal oxidases they use for oxidative phosphorylation (Figure 6). MAG-7, *F. myxofaciens* sp. P3G, and *Ferrovum* sp. JA12 all contain cytochrome *bo*3 type oxidases, but MAG-4 does not contain this type of cytochrome. Similarly, MAGs 4 and 7 and *Ferrovum* JA12 contain a cytochrome *bd*-type quinol oxidase which is not present in *F. myxofaciens* sp. P3G (Figure 6). Terminal oxidases differ in their affinity for oxygen and are differentially expressed depending on the concentration of oxygen [48, 49]. For example, *E. coli* express cytochrome *bo*3 under oxic conditions, and cytochrome *bd*, which has a higher affinity for O_2_, under oxygen limiting conditions [32]. MAG-4 lacks genes for cytochrome *bo*3 which may indicate that this taxon does not colonize well-oxygenated environments that may be more suitable for MAG-7, *F. myxofaciens* sp. P3G, and *Ferrovum* sp. JA12. Similarly, MAGs 4 and 7 and *Ferrovum* sp. JA12 may be able to colonize microaerobic environments that are not ideal for *F. myxofaciens* sp. P3G which lacks high affinity terminal oxidases. In Cabin Branch, these differences may lead to MAGs 4 and 7 preferentially inhabiting different portions of the biofilm with MAG-7 inhabiting the better oxygenated surface of the biofilms and MAG-4 residing in more microaerobic niches within the biofilm.

#### 3.2.2 Nitrogen Cycling

*Ferrovum myxofaciens* sp. P3G can fix nitrogen, but this trait is not conserved within the genus. Neither *Ferrovum* Z-31 nor PN-J185 are capable of nitrogen fixation [15, 18]. At Cabin Branch, MAG-4 encodes the enzymes necessary for N-fixation (NifH, D, and K) and the hydrolysis of urea (Figure 6) while MAG-7 does not. The ability to fix nitrogen would allow MAG-4 to colonize low NH_4_ environments. Additionally, it may influence where MAG-4 resides in the biofilm community as nitrogenase proteins are oxygen sensitive [33]. Therefore, MAG-4 may preferentially reside in low-oxygen environments that provide enough oxygen to perform Fe(II) oxidation, but at low enough concentrations to protect nitrogenase from oxidative damage. This idea is supported by the presence of *cbb*3 and *bd* type oxidases in MAG-4 that are preferentially expressed in microaerobic conditions as discussed above.

#### 3.2.3 Motility

*Ferrovum* spp. vary in their ability to be motile. *Ferrovum* sp. JA-12 and the closely related *Ferrovum* sp. PN-J185 lack genes both for flagella and for chemotaxis whereas *F. myxofaciens* sp. P3G contains the genes necessary for a functional flagellum [9, 15–17, 19]. The MAGs from Cabin Branch are more similar to *F. myxofaciens* sp. P3G in this respect, encoding the genes necessary for the synthesis of a functional flagellum. However, like *F. myxofaciens* sp. P3G, both MAG-4 and MAG-7 lack genes for MotX and MotY which are necessary for flagellar rotation in some species [34]. MAG-7 and *F. myxofaciens* sp. P3G also contain homologs of *tar*, a chemotaxis related protein that responds to the presence of aspartate or maltose and *aer* which produces a protein that allow a chemotactic response to redox state [35, 36] potentially indicating that these taxa can respond to different environmental cues than MAG-4.

## 4. CONCLUSIONS

We recovered co-occurring *Ferrovum* taxa with distinct metabolisms from the emergence and outflow of an AMD site. A DSV-based approach suggested multiple *Ferrovum* taxa co-occur which contrasts to previous OTU-based studies [5]. These data and the *Ferrovum* genomes recovered from metagenomic data highlight the limits of using OTU clustering approaches in identifying discrete species. We also identified the metabolic diversity among closely related *Ferrovum* spp. that likely facilities the co-occurrence of these taxa. Specifically, the differences in nutrient cycling, motility, and chemotaxis may facilitate co-occurrence without direct competition for resources and may also help to explain why *Ferrovum* spp. are nearly ubiquitous in AMD environments despite the geochemical diversity of these environments. These data add to gaps of missing diversity in the *Ferrovum* clade which are ubiquitous in AMD environments but difficult to culture. Finally, our data highlight genetic diversity necessary for closely-related co-occurring species.

## 5. METHODS

### 5.1 Site location and sampling

Cabin Branch is a site in the Daniel Boone National Forest in Kentucky near the border with Tennessee. Limestone was added to the channel as a passive remediation strategy and groundwater flows out from an emergence and across a limestone-lined channel before entering a pond, the Rose Pool (Figure 1). The microbial communities within Cabin Branch are predominantly the Fe(II)-oxidizing taxon Ferrovum myxofaciens [5]. Sample collection was approved by and performed in collaboration with the staff at Daniel Boone National Forest.

### 5.2 Geochemical analyses

#### 5.2.1 Aqueous geochemistry

Temperature and pH were measured onsite using a WTW 330i meter and probe (Xylem Analytics, Weilheim, Germany) and conductivity was measured with a YSI 30 conductivity meter and probe (YSI Inc., Yellow Springs, OH, USA). Total ammonia (NH_4_(T) = NH_3_ + NH_4_^+^) and ferrous iron (Fe(II)) concentration were determined using Hach DR1900 Portable Spectrometers (Hach Company, Loveland, CO) using the salicylate method (Hach Method 8155).

Water samples were collected with a 140 mL syringe directly after rinsing with sample water three times before filling. Water was filtered through 0.8/0.2 μm Supor^®^ syringe filters and dispensed into the appropriate sample bottles following a 10 mL flush to minimize contamination from the filter. Samples for ion chromatography (IC) analysis of major anions (Thermo Scientific Dionex ICS 5000+ ion chromatography system) were stored in 15 mL polypropylene centrifuge tubes. Samples for inductively coupled plasma optical emission spectroscopy (ICP-OES) analysis (Thermo Scientific iCAP 6000 series ICP-OES) of major cations and for inductively coupled plasma mass spectrometry (ICP-MS) analysis (Thermo Scientific X Series 2 ICP-MS) for trace elements were stored in acid-washed (3 day soak in 10 % TraceMetal Grade HNO_3_ (Fisher Scientific, Hampton, NH, USA) followed by triple rinsing with 18.2 MΩ/cm deionized water) 15 mL polypropylene centrifuge tubes and acidified with 400 μL of concentrated, OmniTrace Ultra™ concentrated HNO_3_ (EMD Millipore, Billerica, MA, USA). Field blanks were taken using 18.2 MΩ/cm deionized water transported to the field in 1-liter acid washed Nalgene bottles. IC, ICP-OES, and ICP-MS analyses were carried out by the Analytical Geochemistry Laboratory in the Department of Earth and Environmental Sciences at the University of Minnesota.

Samples for dissolved inorganic carbon (DIC) analysis were filtered into a gas-tight syringe and then injected into Labco Exetainers® (Labco Limited, Lampeter, UK) pre-flushed with He, with excess He removed following introduction of 4 mL of filtered sample with minimal agitation. Samples were stored inverted until and on ice/refrigerated at 4 °C until returned to the lab, where 1 mL of concentrated H_3_PO_4_ was added and the samples shipped to the Stable Isotope Facility (SIF) at the University of California, Davis for analysis. DIC analysis for concentration and ^13^C isotopic signal using a GasBench II system interfaced to a Delta V Plus isotope ratio mass spectrometer (IR-MS) (Thermo Scientific, Bremen, Germany) with raw delta values converted to final using laboratory standards (lithium carbonate, δ^13^C = −46.6 ‰ and a deep seawater, δ^13^C = +0.8 ‰) calibrated against standards NBS-19 and L-SVEC. Samples for dissolved organic carbon (DOC) analysis were filtered through a 0.2 μm polyethersulfone syringe filter that had been flushed with ~ 30 mL of sample and then ~ 40 mL put into a 50 mL centrifuge tube and then immediately flash-frozen on dry ice and kept frozen and in the dark until analysis at the SIF. DOC analysis for concentration and ^13^C isotopic signal were carried out using O.I. Analytical Model 1030 TOC Analyzer (O.I. Analytical, College Station, TX, USA).

#### 5.2.2 CO_2_ assimilation

A microcosm-based approach was employed to assess the potential for inorganic carbon uptake *in situ* through the addition of NaH^13^CO_3_. *In situ* microcosms were performed between noon and 4 PM under full or partial sun. Samples were collected using pre-sterilized spatulas or forceps and ~ 500-mg was placed into pre-combusted (12 h, 450°C) serum vials. Samples were overlaid with spring water from the collected site and serum vials were capped with gas-tight black butyl rubber septa. Assays were initiated by addition of NaH^13^CO_3_ (100 μM final concentration) (Cambridge Isotope Laboratories, Inc., Andover, MA, USA). All assays were performed in triplicate.

We assessed the potential for photoautotrophic (light) and chemoautotrophic (dark) NaH^13^CO_3_ uptake. Assays were stopped by flash freezing vials on dry ice. All vials were stored at −80 °C until processed (described below). Reported values of ^13^C-labelled DIC uptake (carbon fixation rates) reflect the difference in uptake between the biomass in the assays that received NaH^13^CO_3_ and the natural abundance biomass samples described below. For comparisons between mean ^13^C uptake rates, a one-way ANOVA followed by *post hoc* pairwise comparisons between treatments was conducted using a Turkey honest significant difference (HSD) within the R software package (R version 3.3.2). Mean rates with *p*-values < 0.05 were considered significantly different.

#### 5.2.3 C and N concentration and stable isotope signals

Natural abundance samples were collected at the time of sampling, flash frozen on dry ice, stored in liquid N_2_ for transport and stored at −80 °C until processed until processed. Assays were thawed and biomass was washed with HCl (1 M) to remove any extra ^13^C-labelled DIC, triple washed with 18.2 MΩ/cm deionized water, then dried (60°C for three days). Dried biomass was ground/homogenized with a cleaned mortar and pestle (ground with ethanol silica slurry, triple rinsed with 18.2 MΩ/cm deionized water, dried). Dried and ground samples for determination of C and N concentration and stable isotope signal were weighed and placed into tin boats, sealed, and analyzed via an Elementar pyrocube elemental analyzer (EA) periphery connected to an Isoprime 100 continuous flow IRMS (IR-MS) at the University of Minnesota. Linearity corrections were made using NIST Standard 2710, and δ^13^C values were calibrated using reference standards USGS-40 and USGS-41 and checked with a laboratory working standard (glycine). Total uptake of DIC was calculated using DIC numbers determined for the source water (described above). All stable isotope results are given in delta formation expressed as per mil (‰). Carbon stable isotopes are calculated as:

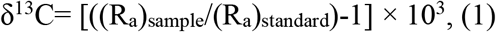

where R_a_ is the ^13^C/^12^C ratio of the sample or standard and are reported versus the Vienna Pee Dee Belemnite (VPDB) standard.

Precautions were taken to minimize cross contamination of natural abundance samples with incubation assays. Natural abundance samples were processed and analyzed in turn, prior to incubation assays. All laboratory processing and weighing equipment was cleaned with 80% ethanol between each sample. Standard checks and blanks were run within each analysis batch (49 samples and standards per batch) to check for memory effects or cross contamination of samples, with none detected.

Assimilation rates were calculated from the following parameters: Absolute isotopic stable carbon ratios were used to determine the difference between the total amount of ^13^C in natural abundance samples and incubation assay replicates. The difference represents total mass of ^13^C-labelled DIC taken up during incubations. Incubation duration times were recorded in the field. Using the organic carbon content, the uptake rate is then calculated from the total μg C taken up divided by the grams of organic C per gram of sediment, and that was divided by the number of hours the incubation was carried out over (typically ~ 2 hours).

### 5.3 Molecular analyses

#### 5.3.1 DNA extraction

DNA was extracted from ~250 mg of biomass using a DNeasy PowerSoil Kit (Qiagen, Carlsbad, CA, USA) according to the manufacturer’s instructions. DNA was extracted from each replicate sample (n=3) and the concentration of DNA was determined using a Qubit dsDNA HS Assay kit and a Qubit 3.0 Fluorometer (Invitrogen, Burlington, ON, Canada). The concentration of DNA in the replicates was within 5% for each sample. Equal volumes of each extraction were pooled, and the concentration of the pooled extract was determined as described above. As a negative DNA extraction control, DNA was extracted from the filter used for the field blank water sample (described above). No DNA was detected in the control and sequencing failed to generate amplicons (see below for amplicon sequencing details).

#### 5.3.2 DNA sequencing

Amplicons were sequenced with MiSeq Illumina 2 × 300 bp chemistry using the primers 515Ff and 806rB targeting V4 hypervariable region of bacterial and archaeal 16S SSU rRNA gene sequences by the University of Minnesota Genomics Center (UMGC). Each sample was sequenced once. For metagenomic sequencing, total DNA was submitted to the UMGC and sequenced using HiSeq2500 High-Output 2 x 125 bp chemistry. The three samples were samples on a single lane.

#### 5.3.3 16S rRNA analysis

16S rRNA sequences from Cabin Branch were used to examine community composition. Primers and unpaired sequences were removed using trimmomatic [37]) The surviving reads were processed in dada2 (v.1.4) following the pipeline tutorial [26]. Briefly, forward reads were trimmed to 210 bp and reverse reads were trimmed to 120 bp based on their quality profiles. Sequences with ambiguous bases and those with more than 2 expected errors were removed. Error rates were estimated using the learnErrors command. Sequences were dereplicated using the derepFastq command and the unique sequence variates were inferred using the dada command. Forward and reverse reads were merged using mergePairs. Contigs shorter than 250 or longer than 256 bp and chimeric sequences were removed. The surviving unique, denoised sequences are referred to as denoised sequence variants (DSVs). Taxonomy was assigned using the Silva training set [38] Eukaryotic sequences and those unclassified at the domain level were removed. The closest cultured and environmental relatives were identified using BLASTN [39].

We retrieved 1,589 aligned, nearly-full length *Ferrovum* sequences from the Silva non-redundant database to serve as a reference alignment for the analyses described below. A 16S rRNA phylogeny was constructed by retrieving the sequences used by Ullrich et al. [18]. These sequences were aligned using MAFFT on the Cipres Science Gateway [40] using the aligned *Ferrovum* sequences from the Silva database as a reference alignment. A tree of the smaller subset of sequences was constructed in RAXML-HPC2 on XSEDE [41] also on the Cipres Science Gateway. Sequences of DSVs classified as *Ferrovum* were added to the alignment using the -addfrags option in MAFFT [42]. Non-full-length sequences were added to the tree using the evolutionary placement algorithm. Trees were rooted and visualized in the interactive tree of life [43]. Raw sequences were uploaded to the NCBI Sequence Read Archive under BioProject PRJNA554371.

#### 5.3.4. Metagenomic analysis

Individual metagenomes were assembled and a co-assembly of all metagenomes was constructed following the “tutorial on assembly-based metagenomics” [44].Trimmed, quality-controlled sequences were assembled using MegaHit [45] using default parameters except minimum contig length, which was set at 1000 base pairs. Reads were mapped to the assembly using bowtie2 [46] and depth was calculated using the jgi_summarize_bam_contig_depths command in Anvi’o [47]. Contigs were binned using default parameters in metabat using [48]. Bin completeness was determined with CheckM [49]. Protein coding regions were identified with prodigal (within CheckM) [50] and GhostKoala was used to annotate protein coding sequences [16, 51].

Each bin was uploaded to KBASE [52] and annotated with Prokka using the “Annotate Assembly and Re-annotate Genomes with Prokka (v1.12)” app. A concatenated gene tree containing each bin and four published *Ferrovum* spp. genomes was constructed using the “Insert Set of Genomes Into Species Tree 2.1.10” app. MAGs that were more closely related to *Ferrovum* spp. than other taxa were selected for further analysis. The pairwise average nucleotide identity (ANI) between each bin and published *Ferrovum* genomes was calculated using anvi-compute-ani in Anvi’o. If MAGs shared >98% pairwise ANI and were phylogenetically cohesive, we considered them to be representing the same populations and the bin with the highest completeness was chosen for further analysis. Raw reads, assemblies, and MAGs were uploaded to the NCBI Sequence Read Archive under BioProject PRJNA554371.

##### 16S rRNA

A custom blast database containing the 16S rRNA gene from the *Ferrovum myxofaciens* type strain (NR_117782.1) using the makeblastdb command in BLAST+. Each of the *Ferrovum* bins were searched for 16S rRNA genes using BLAST and a cutoff value of E-30.

##### Cyc-2

A custom blast database of the high molecular weight Cyc-2 like proteins was constructed using the methods described above. Each of the *Ferrovum* bins were searched for the Cyc-2-like protein identified by Ullrich et al.,[16] by blasting translated nucleotide sequences against the blast database with an E-value of E-120. The retrieved sequences, database sequences, and a sequence for Cyc-2 from an *Acidithiobacillus sp*. (AVK42811.1) were aligned using MAFFT and a maximum likelihood tree was constructed as described above. The tree was rooted on *Acidithiobacillus* and visualized in iTOL.

#### 5.3.5. Comparison with other Ferrovum genomes

Existing *Ferrovum* genomes are located within either the Group I or Group IV *Ferrovum* clades [18]. The genome for *Ferrovum* sp. JA12 (Group I) and *Ferrovum myxofaciens* (Group IV) were downloaded from NCBI. CheckM was used to determine genome completeness, and within CheckM, prodigal was used to identify protein coding sequences. Ghost KOALA was used to annotate protein coding sequences using the genus_prokaryotes database. Protein coding sequences were also annotated with prokka. We then manually compared the gene content of MAGs to that of existing *Ferrovum* genomes.

## ABBREVIATIONS

AMD: Acid mine drainage
ANI: Average nucleotide identity
APS: Ammonium persulfate 3-phosphoadenosine-5-phosphosulfate
DIC: Dissolved inorganic carbon
DSV: Denoised sequence variants
ICP-MS: Inductively coupled plasma mass spectrometry
ICP-OES: Inductively coupled plasma atomic emission spectroscopy
MAG: Metagenome assembled genomes
PAPS: 3-phosphoadenosine-5-phosphosulfate
TCA: Tricarboxylic acid cycle

## ACKNOWLEDGEMENTS

We are grateful to the staff of the National Forest Service and Daniel Boone National Forest, especially Margueritte Wilson and Claudia Cotton, for the advice and insight regarding mine locations. We thank A. Gangidine, M. Berberich, R. Jain, and C. Schuler for assistance in field sampling and processing samples in the laboratory.

## AVAILABILITY OF DATA AND MATERIALS

All analyses tools used in the study are publicly available. 16S rRNA and functional genes of closely related species and outgroups were downloaded from NCBI databases and accession numbers are provided for these sequences. Raw reads and assembled scaffolds for the metagenomes and MAG sequences as well as the amplicon libraries have been deposited in the NCBI under BioProject PRJNA554371.

## AUTHORS’ CONTRIBUTIONS

Drs. Havig, Grettenberger, and Hamilton contributed to the conception of the project, data analysis interpretation of results, creation of figures, writing and editing the manuscript. Drs. Havig and Hamilton performed sample collection and sample analysis.

## ETHICS APPROVAL AND CONSENT TO PARTICIPATE

Not applicable.

## CONSENT FOR PUBLICATION

Not applicable.

## REFERENCES

1. Baker BJ, Banfield JF. Microbial communities in acid mine drainage. Fems Microbiol Ecol. 2003;44:139–52.

2. Johnson BD. Chemical and Microbiological Characteristics of Mineral Spoils and Drainage Waters at Abandoned Coal and Metal Mines. Water Air Soil Pollut Focus. 2003;3:47–66.

3. Schippers A, Breuker A, Blazejak A, Bosecker K, Kock D, Wright TL. The biogeochemistry and microbiology of sulfidic mine waste and bioleaching dumps and heaps, and novel Fe(II)-oxidizing bacteria. Hydrometallurgy. 2010;104:342–50.

4. Grettenberger CL, Pearce AR, Bibby KJ, Jones DS, Burgos WD, Macalady JL. Efficient Low-pH Iron Removal by a Microbial Iron Oxide Mound Ecosystem at Scalp Level Run. Appl Environ Microb. 2017;83:e00015–17.

5. Havig JR, Grettenberger C, Hamilton TL. Geochemistry and microbial community composition across a range of acid mine drainage impact and implications for the Neoarchean‐ Paleoproterozoic transition. J Geophys Res Biogeosciences. 2017;122:1404–22.

6. Jones DS, Kohl C, Grettenberger C, Larson LN, Burgos WD, Macalady JL. Geochemical Niches of Iron-Oxidizing Acidophiles in Acidic Coal Mine Drainage. Appl Environ Microb. 2015;81:1242–50.

7. Johnson BD, Hallberg KB, Hedrich S. Uncovering a Microbial Enigma: Isolation and Characterization of the Streamer-Generating, Iron-Oxidizing, Acidophilic Bacterium “Ferrovum myxofaciens.” Appl Environ Microb. 2014;80:672–80.

8. Hallberg KB, Coupland K, Kimura S, Johnson BD. Macroscopic Streamer Growths in Acidic, Metal-Rich Mine Waters in North Wales Consist of Novel and Remarkably Simple Bacterial Communities. Appl Environ Microb. 2006;72:2022–30.

9. Schlömann M, Kipry J, Mosler S, Poehlein A, Keller A, Janneck E, et al. Physiological, Genomic, and Proteomic Characterization of New “Ferrovum” Strains Obtained from a Pilot Plant for Mine-Water Treatment. Adv Mat Res. 2013;825:149–52.

10. Johnson DB. Acidophilic microbial communities: Candidates for bioremediation of acidic mine effluents. Int Biodeter Biodegr. 1995;35:41–58.

11. Kipry J, Jwair R, Gelhaar N, Wiacek C, Janneck E, Schlömann M. Enrichment of “Ferrovum” spp. and Gallionella Relatives Using Artificial Mine Water. Adv Mat Res. 2013;825:54–7.

12. Tischler JS, Jwair R, Gelhaar N, Drechsel A, Skirl A-M, Wiacek C, et al. New cultivation medium for “Ferrovum” and Gallionella-related strains. J Microbiol Meth. 2013;95:138–44.

13. Kuang J-L, Huang L-N, Chen L-X, Hua Z-S, Li S-J, Hu M, et al. Contemporary environmental variation determines microbial diversity patterns in acid mine drainage. Isme J. 2013;7:1038.

14. Kay C, Rowe O, Rocchetti L, Coupland K, Hallberg K, Johnson D. Evolution of Microbial “Streamer” Growths in an Acidic, Metal-Contaminated Stream Draining an Abandoned Underground Copper Mine. Life. 2013;3:189–210.

15. Ullrich SR, González C, Poehlein A, Tischler JS, Daniel R, Schlömann M, et al. Gene Loss and Horizontal Gene Transfer Contributed to the Genome Evolution of the Extreme Acidophile “Ferrovum.” Front Microbiol. 2016;7:797.

16. Ullrich SR, Poehlein A, Tischler JS, González C, Ossandon FJ, Daniel R, et al. Genome Analysis of the Biotechnologically Relevant Acidophilic Iron Oxidising Strain JA12 Indicates Phylogenetic and Metabolic Diversity within the Novel Genus “Ferrovum.” Plos One. 2016;11:e0146832.

17. Mosler S, Poehlein A, Voget S, Daniel R, Kipry J, Schlömann M, et al. Predicting the Metabolic Potential of the Novel Iron Oxidising Bacterium “Ferrovum” sp. JA12 Using Comparative Genomics. Adv Mat Res. 2013;825:153–6.

18. Ullrich SR, Poehlein A, Daniel R, Tischler JS, Vogel S, Schlömann M, et al. Comparative Genomics Underlines the Functional and Taxonomic Diversity of Novel “*Ferrovum*” Related Iron Oxidizing Bacteria. Adv Mat Res. 2015;1130:15–8.

19. Moya-Beltrán A, Cárdenas J, Covarrubias PC, Issotta F, Ossandon FJ, Grail BM, et al. Draft Genome Sequence of the Nominated Type Strain of “Ferrovum myxofaciens,” an Acidophilic, Iron-Oxidizing Betaproteobacterium. Genome Announc. 2014;2:e00834–14.

20. Bowers RM, Kyrpides NC, Stepanauskas R, Harmon-Smith M, Doud D, Reddy T, et al. Minimum information about a single amplified genome (MISAG) and a metagenome-assembled genome (MIMAG) of bacteria and archaea. Nat Biotechnol. 2017;35:725.

21. Peña A, Teeling H, Huerta-Cepas J, Santos F, Yarza P, Brito-Echeverría J, et al. Fine-scale evolution: genomic, phenotypic and ecological differentiation in two coexisting Salinibacter ruber strains. Isme J. 2010;4:882.

22. Antón J, Lucio M, Peña A, Cifuentes A, Brito-Echeverría J, Moritz F, et al. High Metabolomic Microdiversity within Co-Occurring Isolates of the Extremely Halophilic Bacterium Salinibacter ruber. Plos One. 2013;8:e64701.

23. Meyer JL, Huber JA. Strain-level genomic variation in natural populations of Lebetimonas from an erupting deep-sea volcano. Isme J. 2014;8:867.

24. Rodriguez-R LM, Castro JC, Kyrpides NC, Cole JR, Tiedje JM, Konstantinidis KT. How Much Do rRNA Gene Surveys Underestimate Extant Bacterial Diversity? Appl Environ Microb. 2018;84:e00014–18.

25. Callahan BJ, McMurdie PJ, Holmes SP. Exact sequence variants should replace operational taxonomic units in marker-gene data analysis. Isme J. 2017;11:ismej2017119.

26. Callahan BJ, McMurdie PJ, Rosen MJ, Han AW, Johnson AA, Holmes SP. DADA2: High-resolution sample inference from Illumina amplicon data. Nat Methods. 2016;13:nmeth.3869.

27. Figueras M, Beaz-Hidalgo R, Hossain MJ, Liles MR. Taxonomic Affiliation of New Genomes Should Be Verified Using Average Nucleotide Identity and Multilocus Phylogenetic Analysis. Genome Announc. 2014;2:e00927–14.

28. MacArthur RH. Population Ecology of Some Warblers of Northeastern Coniferous Forests. Ecology. 1958;39:599–619.

29. Inouye DW. Resource Partitioning in Bumblebees: Experimental Studies of Foraging Behavior. Ecology. 1978;59:672–8.

30. Swanson BO, Gibb AC, Marks JC, Hendrickson DA. Trophic Polymorphism And Behavioral Differences Decrease Intraspecific Competition In A Cichlid, Herichthys Minckleyi. Ecology. 2003;84:1441–6.

31. Chan C, McAllister SM, Garber A, Hallahan BJ, Rozovsky S. Fe oxidation by a fused cytochrome-porin common to diverse Fe-oxidizing bacteria. Biorxiv. 2018;:228056.

32. Cotter P, Chepuri V, Gennis R, Gunsalus R. Cytochrome o (cyoABCDE) and d (cydAB) oxidase gene expression in Escherichia coli is regulated by oxygen, pH, and the fnr gene product. J Bacteriol. 1990;172:6333–8.

33. Gallon JR. The oxygen sensitivity of nitrogenase: a problem for biochemists and micro-organisms. Trends Biochem Sci. 1981;6:19–23.

34. Koerdt A, Paulick A, Mock M, Jost K, Thormann KM. MotX and MotY Are Required for Flagellar Rotation in Shewanella oneidensis MR-1▿ †. J Bacteriol. 2009;191:5085–93.

35. Rebbapragada A, Johnson MS, Harding GP, Zuccarelli AJ, Fletcher HM, Zhulin IB, et al. The Aer protein and the serine chemoreceptor Tsr independently sense intracellular energy levels and transduce oxygen, redox, and energy signals for Escherichia coli behavior. Proc National Acad Sci. 1997;94:10541–6.

36. Bibikov S, Biran R, Rudd K, Parkinson J. A signal transducer for aerotaxis in Escherichia coli. J Bacteriol. 1997;179:4075–9.

37. Bolger AM, Lohse M, Usadel B. Trimmomatic: a flexible trimmer for Illumina sequence data. Bioinformatics. 2014;30:2114–20.

38. Quast C, Pruesse E, Yilmaz P, Gerken J, Schweer T, Yarza P, et al. The SILVA ribosomal RNA gene database project: improved data processing and web-based tools. Nucleic Acids Res. 2013;41:D590–6.

39. Camacho C, Coulouris G, Avagyan V, Ma N, Papadopoulos J, Bealer K, et al. BLAST+: architecture and applications. Bmc Bioinformatics. 2009;10:421.

40. Miller MA, Pfeiffer W, Schwartz T. Creating the CIPRES Science Gateway for Inference of Large Phylogenetic Trees. 2010 Gatew Comput Environ Work Gce. 2010;:1–8.

41. Stamatakis A. RAxML-VI-HPC: maximum likelihood-based phylogenetic analyses with thousands of taxa and mixed models. Bioinformatics. 2006;22:2688–90.

42. Katoh K, Standley DM. MAFFT Multiple Sequence Alignment Software Version 7: Improvements in Performance and Usability. Mol Biol Evol. 2013;30:772–80.

43. Letunic I, Bork P. Interactive tree of life (iTOL) v3: an online tool for the display and annotation of phylogenetic and other trees. Nucleic Acids Res. 2016;44:W242–5.

44. Eren M. A tutorial on assembly-based metagenomics.

45. Li D, Liu C-M, Luo R, Sadakane K, Lam T-W. MEGAHIT: an ultra-fast single-node solution for large and complex metagenomics assembly via succinct de Bruijn graph. Bioinformatics. 2015;31:1674–6.

46. Langmead B, Salzberg SL. Fast gapped-read alignment with Bowtie 2. Nat Methods. 2012;9:357.

47. Eren MA, Esen ÖC, Quince C, Vineis JH, Morrison HG, Sogin ML, et al. Anvi’o: an advanced analysis and visualization platform for ‘omics data. Peerj. 2015;3:e1319.

48. Kang DD, Froula J, Egan R, Wang Z. MetaBAT, an efficient tool for accurately reconstructing single genomes from complex microbial communities. Peerj. 2015;3:e1165.

49. Parks DH, Imelfort M, Skennerton CT, Hugenholtz P, Tyson GW. CheckM: assessing the quality of microbial genomes recovered from isolates, single cells, and metagenomes. Genome Res. 2015;25:1043–55.

50. Hyatt D, Chen G-L, LoCascio PF, Land ML, Larimer FW, Hauser LJ. Prodigal: prokaryotic gene recognition and translation initiation site identification. Bmc Bioinformatics. 2010;11:119.

51. Kanehisa M, Sato Y, Morishima K. BlastKOALA and GhostKOALA: KEGG Tools for Functional Characterization of Genome and Metagenome Sequences. J Mol Biol. 2016;428:726–31.

52. Arkin AP, Cottingham RW, Henry CS, Harris NL, Stevens RL, Maslov S, et al. KBase: The United States Department of Energy Systems Biology Knowledgebase. Nat Biotechnol. 2018;36:566.

