## Supplemental Table 1 for "Intra-genus metabolic diversity facilitates co-occurrence of multiple *Ferrovum* species at an acid mine drainage site"

Supplementary Table 1. Carbon uptake experiment results.

| Site and Biofilm Description | Treatment | Time | Carbon uptake experiments | |
|---|---|---|---|---|
| | | | Average | S.D. |
| Emergence | | (hrs) | (μg C uptake/g $C_{biomass}$/hr) | |
| Green/White Biofilm | Light | 1 | 32.55 | 7.95 |
| Green only | Dark | 1 | 1.63 | 0.45 |
| White only | Dark | 1 | 0.31 | 0.40 |
| Green/White Biofilm | Light | 2 | 21.69 | 6.36 |
| Green only | Dark | 2 | 0.75 | 0.17 |
| White only | Dark | 2 | 0.66 | 0.24 |
| Outflow 1 | | | | |
| Yellow/Green Biofilm | Light | 1 | 35.95 | 5.52 |
| Yellow/Green Biofilm | Dark | 1 | 2.59 | 0.15 |
| Yellow/Green Biofilm | Light | 2 | 15.09 | 0.11 |
| Yellow/Green Biofilm | Dark | 2 | 1.25 | 0.03 |
| Outflow 2 | | | | |
| Rose Pool Sediments | Light | 2 | 1.37 | 0.06 |
| Rose Pool Sediments | Dark | 2 | 0.07 | 0.13 |

Average = average of three replicates, S.D. = standard deviation.
